# Temporal variation in lymphocyte proteomics

**DOI:** 10.1101/2021.07.29.454362

**Authors:** Michaela A. McCown, Carolyn Allen, Daniel D. Machado, Hannah Boekweg, Yiran Liang, Andikan J. Nwosu, Ryan T. Kelly, Samuel H. Payne

**Affiliations:** Biology Department, Brigham Young University, Provo UT 84604; Chemistry and Biochemistry Department, Brigham Young University, Provo, UT 84604

## Abstract

Chronic Lymphocytic Leukemia (CLL) is a slow progressing disease, characterized by a long asymptomatic stage followed by a symptomatic stage during which patients receive treatment. While proteomic studies have discovered differential pathways in CLL, the proteomic evolution of CLL during the asymptomatic stage has not been studied. In this pilot study, we show that by using small sample sizes comprising ~145 cells, we can detect important features of CLL necessary for studying tumor evolution. Our small samples are collected at two time points and reveal large proteomic changes in healthy individuals over time. A meta-analysis of two CLL proteomic papers showed little commonality in differentially expressed proteins and demonstrates the need for larger control populations sampled over time. To account for proteomic variability between time points and individuals, large control populations sampled at multiple time points are necessary for understanding CLL progression. Data is available via ProteomeXchange with identifier PXD027429.

## Introduction

Chronic Lymphocytic Leukemia (CLL) is the most common blood plasma cancer for individuals over the age of 65. There are an estimated 21,000 new cases and 4,000 deaths in the United States each year^1^. About 80% of new cases are diagnosed in a pre-symptomatic early stage, where therapies have been shown to be ineffective and therefore are not administered^2^. This stage is followed by a symptomatic stage with a variety of chemotherapy options^3^. Immunotherapy with CAR T-cell treatment has also been attempted with limited success^4,5^. Treatments generally delay terminal illness but do not cure the disease. To find more effective treatment options, research into long term disease evolution is essential.

Studies in the fields of genomics and transcriptomics have yielded insight into the characterization, diagnosis, and treatment of CLL. Genetic and genomic profiling has identified a variety of mutations common in CLL that help predict disease progression. Mutations in the immunoglobulin heavy-chain variable region gene (*IGHV*) are correlated with better outcomes^6,7^, while mutations in NOTCH1, SF3B1, and BIRC3 are all correlated with poorer outcomes^8^. Transcriptomic profiling has been used to identify predisposition to the disease, severity, and time to treatment^9–11^. Relatively few studies have explicitly looked at longitudinal evolution of CLL; however, one recent genomic study showed that when sampled at multiple time points, many CLL patients have minimal genetic subclonal evolution during early disease development^12^.

Proteomics provides unique insight into the characteristics of CLL by identifying differentially expressed proteins and pathways. Johnston et al. compared the B cells of healthy donors to CLL patients^13^. They found several surface proteins that contribute to CLL development and may be used as therapeutic targets. Mayer et al. investigated proteomic distinctions between B cells from young healthy controls, elderly healthy controls, and CLL patients^14^. They discovered that the proteome of elderly B-cells is closely related to leukemia, indicating that elderly individuals are at a greater risk of developing CLL. Single-cell proteomics has the potential to increase our understanding of tumor composition. In a related leukemia, Acute Myeloid Leukemia, single-cell proteomics has successfully demonstrated the possibility of measuring inter-cellular heterogeneity^15^.

While the proteomic differences between CLL and healthy B lymphocytes have been characterized, the temporal changes during disease progression and cellular heterogeneity have not been explored. For any disease, understanding changes across the natural history provides important insight for developing improved treatment options and identifying prognostic markers. Since CLL has a long asymptomatic developmental phase, this evolution is even more vital to understand. However, longitudinal CLL studies are difficult to perform due to the size of samples required for current proteomic experiments. Small sample sizes will allow for samples to be taken more frequently and to characterize rare cell types. In this pilot study, we characterize the proteome of lymphocytes from two healthy subjects at two time points using ultra-small samples. We demonstrate that the lymphocyte proteome in healthy individuals fluctuates widely with time and between individuals. We also perform a meta-analysis with other proteomics studies and find that our ultra-small proteomics sampling has substantial overlap with these traditional bulk sample studies. This demonstrates our samples have sufficient coverage depth for longitudinal study of CLL.

## Methods

### Lymphocyte Collection

At each sampling time, whole blood samples were collected from the Brigham Young University’s Student Health Center (IRB Authorization X19045: Understanding the Progression of Chronic Lymphocytic Leukemia). Samples were taken from two healthy adult male volunteers. Samples were kept on ice for transport and prior to processing. These samples were collected in June and July of 2020, thirty days apart.

Sample preparation is outlined and archived on Protocols.io^16^. Cells were processed in triplicate from 200 μl whole blood from a 4ml EDTA tube. White blood cells were isolated by lysing the red blood cells and pelleting the intact cells. 200 μl of whole blood was mixed with 4 ml of red blood cell lysis buffer in 5 ml falcon tubes and incubated for 5 minutes at room temperature. Debris was separated from intact white blood cells by centrifugation at 250 ×g for 5 minutes and the supernatant was decanted. In addition to the three replicates for each subject, we also prepared four replicates for FACS controls (cells only, APC-positive, FITC-positive, and PI-positive), which started with 100 μl of whole blood. Since the PI-positive control is designed to mark dead or dying cells by staining exposed DNA, this control sample used dead cells from month-old refrigerated blood.

B and T cells were isolated by immunostaining and FACS sorting. T lymphocytes and B lymphocytes were marked using CD3 anti-human FITC and CD19 anti-human APC (BioLegend FITC: 300305, APC: 392503), respectively. Cells were resuspended in 100 μl of phosphate buffer saline (PBS) with 5 μl of each immunostain and 10 μl PI. Stain was added to the appropriate single color controls. Samples were incubated for 20 minutes in the dark at room temperature. Excess dye was removed by washing with 2 ml PBS, then centrifuging at 250 ×g for 5 minutes and decanting the supernatant. The cells were resuspended in 300 μl PBS. The cells were filtered through 40 μm filters, bringing the cell suspension to an appropriate concentration for FACS sorting. B cells and T cells were then FACS sorted to samples of 145 cells in 384 well plates that were prewashed with water, mobile phase A, 0.01% n-dodecyl β-D-maltoside (DDM) in stages and prefilled with 4 μl of 0.1% DDM to minimize cell damage from sorting.

### Protein Quantification by Mass Spectrometry

Samples were analyzed using global proteomics for low input samples. First, protein was extracted using automated digestion and nanoLC-MS/MS as described by Liang et al^17^. Cells were sonicated with 4 μl of 0.1% DDM (n-dodecyl β-D-maltoside) for 5 minutes and centrifuged at 1500 rpm for 1 minute after sorting. The OT-2 liquid handler (Opentrons, Brooklyn, NY) carried out further sample processing as reported previously^17^. In brief, the liquid handler added the following: 1 μl of 25 mM dithiothreitol, then 2 μl of 50mM ABC buffer every 15 minutes during a 1-hour incubation at 70 □, 1 μl of 50 mM iodoacetamide prior to a 30 minute incubation at 25□, 1 μl of 1 ng/μl lys-C prior to a 3 hour incubation at 37□, 1 μl of 1ng/μl trypsin, 2 μl of water every 3 hours during a 12 hour incubation at 37□, 1 μl of water and 1 μl of 5% formic acid prior to a 1 hour incubation at 25□. At that time, samples were stored at −20□ until thawing for nanoLC-MS/MS.

Five replicates of each experimental condition (cell type, subject, time point) were run on an Orbitrap Exploris 480 (Thermo Fisher) mass spectrometer using a Nanospray Flex ion source at 2.0 kV. For additional settings of the LC-MS/MS data acquisition, see the methodological details in reference^17^. The mass spectrometry proteomics data have been deposited to the ProteomeXchange Consortium^18^ via the PRIDE^19^ partner repository with the dataset identifier PXD027429. Raw mass spectrometry files were processed using the FragPipe interface with MSFragger^20^ and IonQuant^21^ using default settings. Match Between Runs was not used. False discovery rates are <0.01. The database search was conducted with a UniProt human FASTA file (Swiss-Prot, reviewed, and downloaded July 2020).

### Relationship Analysis

Specific analyses were conducted in Python using Jupyter Notebooks. Full code can be found on GitHub at https://github.com/PayneLab/TwoMonthLymphocyte following a standard analysis sharing pattern^22^. The quantification tables were read in as Pandas dataframes from the longitudinalCLL package. Quantification tables were then normalized by log 2 transformation. Because batch effects shift the signal intensity between runs, each run was adjusted by its median value to create a normalized dataset of relative intensities centered at zero. Identifications by sample are counted by the number of protein groups with a non-null intensity, i.e., those that were measured at any abundance. Gene set enrichment analysis^23,24^ was done using the cell type classifier ProteomicsDB^25^ to verify these proteins as lymphocyte-representative expressions. Proteins were filtered by cell type for those that are present in at least half of the samples. Spearman correlation coefficients were calculated to show reproducibility within a cell type. To better display the change between B lymphocytes collected at each time, the fold change was calculated for each protein and graphed on a log 2 scale. These figures were created using Seaborn and pyplot. Significance of relationships was determined by t-test for independent samples with Bonferroni correction. Proteins with a p-value <.05 were considered significant. KEGG pathway analysis was done using gprofiler (https://biit.cs.ut.ee/gprofiler) to determine the relationships between cell types, subjects, and time points. The full software for analysis and figure creation is found in our GitHub repository, specifically see ~/Make_Figure_1A.ipynb, ~/Make_Figure_1B.ipynb, ~/Make_Figure_1C.ipynb, and ~/Make_Figure_1D.ipynb.

### Cross Study Analysis

Data from two other recent proteomics studies on CLL was analyzed. Each study isolated proteins from CLL and control subjects, analyzed them by MS/MS, and identified differentially expressed proteins. Each study used slightly different methods at each step, which we review here briefly; for a more comprehensive comparison, see Supplementary Table 3. The full software for analysis and figure creation is found in our GitHub repository, specifically see ~/Mayer_Abundance_Comp.ipynb, ~/Make_Figure_2.ipynb and ~/Make_Figure_3.ipynb. For table creation, see ~/Make_Supplementary_Table_1.ipynb, ~/Make_Supplementary_Table_2, and ~/Make_Supplementary_Table_4.ipynb.

Mayer et al.^14^ compared proteome data from 9 CLL patients to 3 age-matched and 3 young healthy controls, noting that changes correlated with aging mirror the changes correlated with CLL. B lymphocytes were extracted via Ficoll Paque centrifugation and anti-CD19 magnetic beads for all samples. 20 μg of protein was analyzed using a Q Exactive Orbitrap. Data was analyzed using MaxQuant and identified 6,945 unique proteins. Protein abundance between CLL and age-matched controls were compared by t test, with significantly altered proteins those with FDR<0.05 following a permutation-based multiparameter correction. Data was published as Table S3.

Johnston et al.^13^ compared proteome data from 14 CLL patients to 3 healthy donors, highlighting altered pathways that are targeted by existing therapeutics. PMBCs were isolated by density gradient centrifugation. In the CLL patients, these cells were >90% CD19+/CD3+ so no further separation was required. □500 ng of peptides was analyzed using an LQT-Orbitrap Elite Velos Pro hybrid. Data was analyzed using Proteome Discoverer and identified 8,694 unique proteins, 5,956 of which were quantified. Protein abundance was compared by t test on the log2 ratios between CLL patients and each healthy control, with significantly altered proteins those with p< 0.05.

For each of these studies, data was downloaded from the respective supplementary table and compared in a Jupyter Notebook. The lists of all identified proteins and differentially expressed proteins were compared for proteins identified in multiple studies. KEGG pathway analysis was done using gprofiler (https://biit.cs.ut.ee/gprofiler) to show whether the proteins identified as differentially expressed represent the same pathways in each study.

## Results

Getting high quality data from ultra-small samples at multiple time points is vital to understanding the evolution of a heterogeneous and dynamic disease like CLL. As a pilot study to test the technological capabilities for ultra-small sample longitudinal analysis, we characterized the global proteomics of 145-cell B cell and T cell samples from two healthy subjects at two time points (see Methods).

### B cell variability

Global proteomic data for FACS isolated B cells and T cells from two healthy subjects at two time points identified 2,426 unique proteins (Supplementary Table 1). Five technical replicates were run for each subject and cell type at each time point. The data had high technical reproducibility as shown by an average Spearman correlation coefficient of 0.93 between sample replicates. When comparing subjects, we found 91 proteins upregulated in subject 1, and 130 proteins upregulated in subject 2 (Figure 1A). A KEGG pathway enrichment of differential proteins showed no substantial difference in pathways between the two subjects. When comparing B and T cells in subject 1, we found 131 up-regulated proteins in B cells, and 182 up-regulated in T cells (Figure 1B). A KEGG pathway enrichment showed expected enrichment of pathways related to each cell’s characteristic function (Supplementary Table 2). For example, B cell receptor signaling was elevated in B cells, while leukocyte transendothelial migration was elevated in T cells.

**Figure 1.**
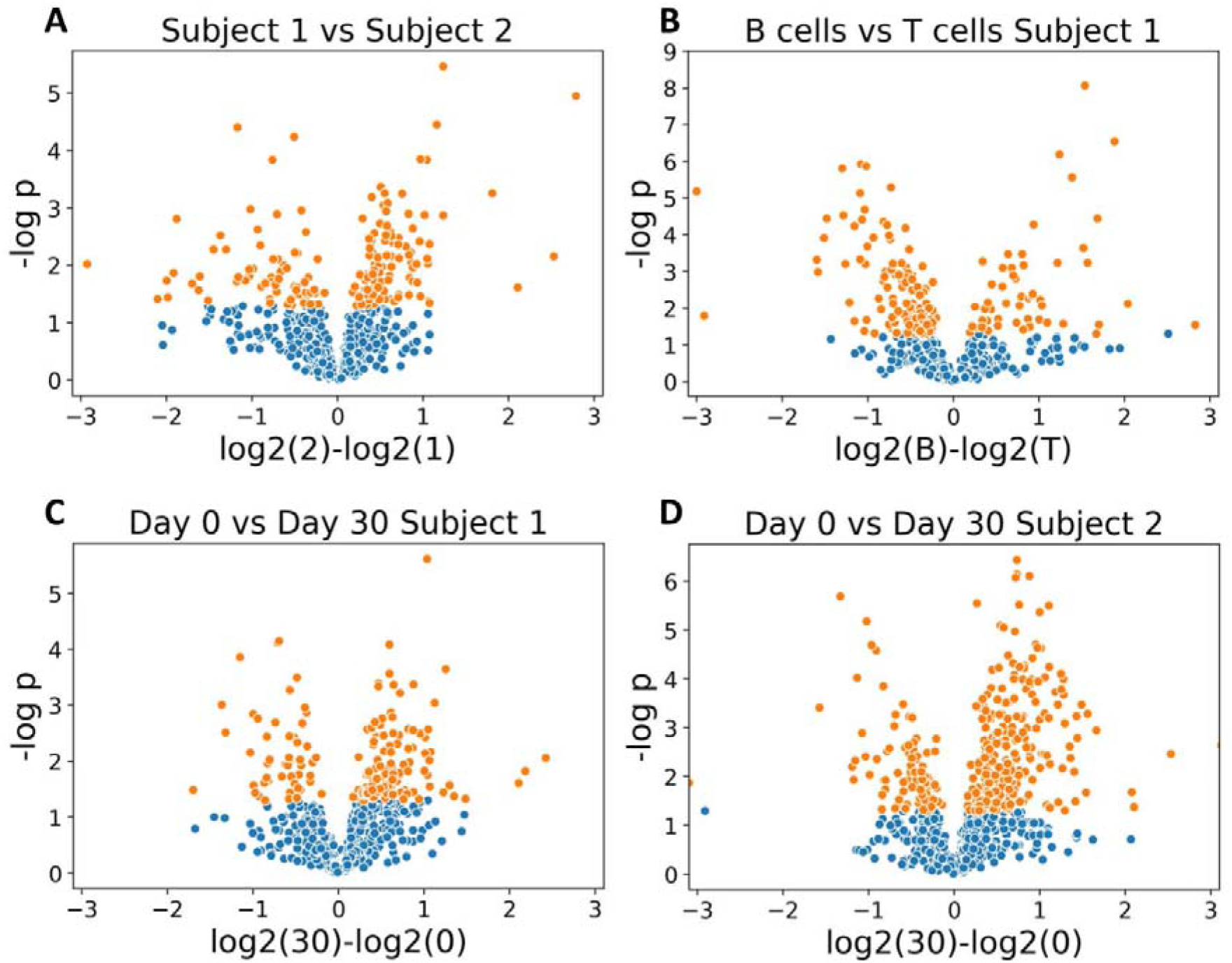
A) Volcano plot of t tests between proteins found in subject 1 B cells and subject 2 B cells. Orange data points indicate differential proteins. B) Volcano plot of t tests between proteins found in B cells and T cells for subject 1. C, D) Volcano plot of t tests between proteins found in June and July for subject 1 and subject 2, respectively. Within our GitHub repository (https://github.com/PayneLab/TwoMonthLymphocyte), the exact code for Figure 1 can be found in files “~/Make_Figure_1A.ipynb”, “~/Make_Figure_1B.ipynb”, “~/Make_Figure_1C.ipynb”, “~/Make_Figure_1D.ipynb”

Molecular datasets used to identify differences between disease and control individuals frequently only include a single time point, excluding the characterization of natural variability that occurs within an individual over time. We performed a variety of comparisons between experimental conditions to identify temporal proteomic changes. We found significant changes in the proteins identified at day 0 compared to those identified at day 30 for subject 1, with 62 upregulated proteins in June, and 196 upregulated in July (Figure 1C). Subject 2 experienced similar changes between time points, with 92 upregulated proteins in June, and 440 upregulated in July (Figure 1D). A pathway analysis based on time differences showed no major difference in enriched pathways for either subject. The change between time points is similar to the differences between subjects and between cell types, both in terms of the number of differential proteins and also the magnitude of change. However, these temporal protein changes seem to be random protein fluctuations as we identified altered pathways when comparing B cells and T cells, but no substantially altered pathways in the time point and subject comparisons.

### Meta-analysis of CLL proteomics

Since we wanted to determine whether our ultra-small samples identified key features which would be useful in the study of healthy and leukemic B cells, we evaluated how our protein identifications compared to two published CLL studies. Two studies compared global proteomic data from healthy donor B cells to B-CLL cells at one time point13,14. Both studies had similar methods for sample preparation, data acquisition and analysis (Supplementary Table 3). Both studies had three control samples and similar numbers of subjects. Importantly, the supplemental data provided in both papers allowed us to perform a meta-analysis and identify commonalities which distinguish healthy and leukemic B cells. Johnston et al. quantified 5,956 proteins and Mayer et al. quantified 6,945 unique proteins. Of these proteins, 5,328 proteins were identified by both studies (Figure 2A).

**Figure 2.**
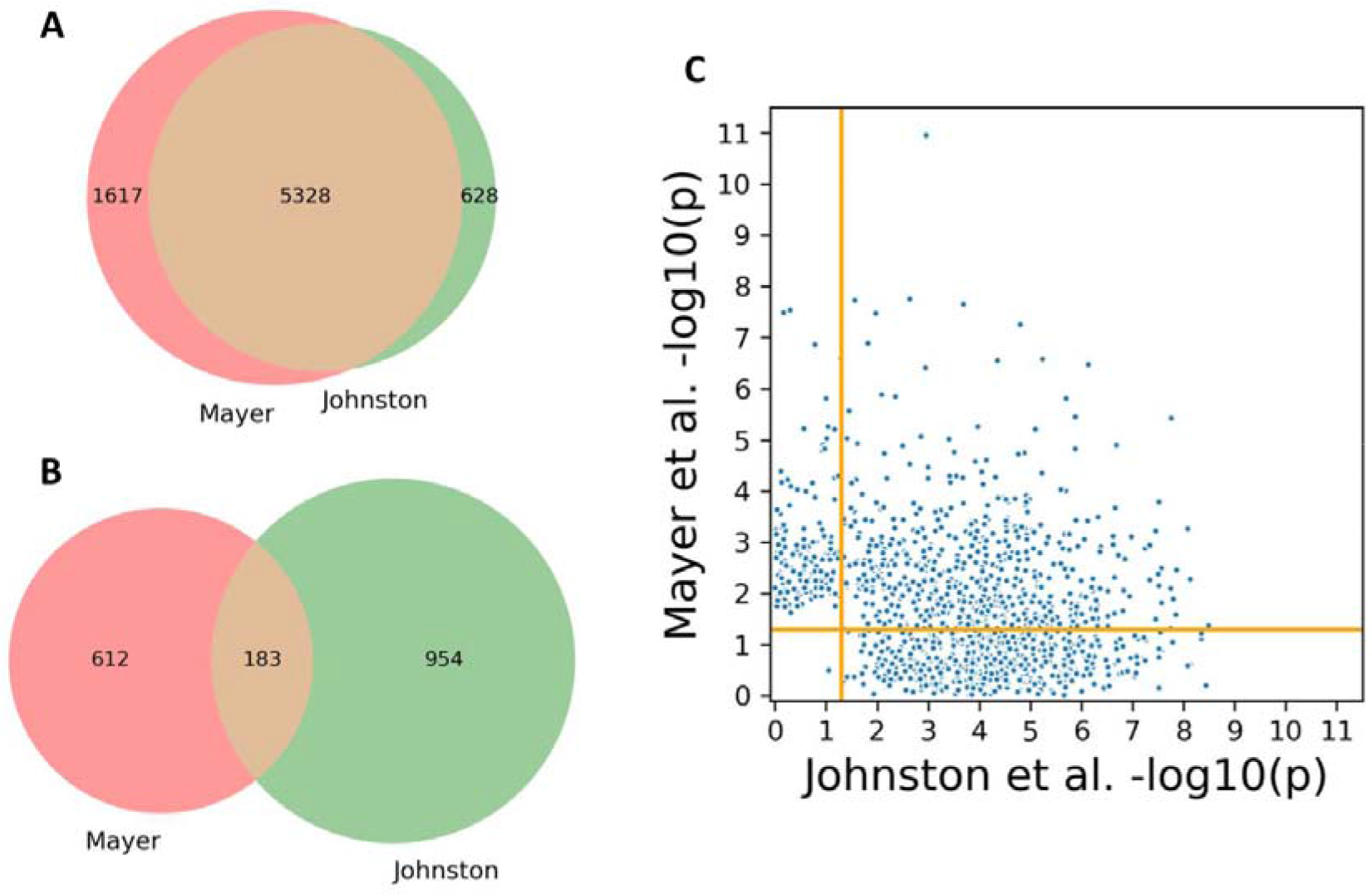
A) Venn diagram of total proteomic data from Mayer et al.^14^ and Johnston et al.^13^ B) Venn diagram of proteins Mayer et al.^14^ and Johnston et al.^13^ identified as differential. C) Scatterplot of p-values for proteins Mayer et al.^14^ and Johnston et al.^13^ identified as differential (r = −0.27). Within our GitHub repository (https://github.com/PayneLab/TwoMonthLymphocyte), the exact code for Figure 2 can be found in file “~/Make_Figure_2.ipynb”

Each study also listed the differentially expressed proteins between healthy donor B cells and B-CLL cells. To determine which proteins were differentially expressed, Mayer et al.^14^ used a t test (p < 0.05), while Johnston et al.^13^ used regulation score and ran the t test on minimal log ratios. Mayer et al.^14^ identified 795 differentially expressed proteins and Johnston et al.^13^ identified 1137. Of these proteins, only 183 proteins were identified as differentially expressed in both studies, demonstrating that the studies identified substantially different sets of differentially expressed proteins (Figure 2B). To determine the relationship between proteins considered significant in one study but not in another we compared the protein p-values from Mayer et al.s to those of Johnston et al. (Figure 2C). The comparison had a Pearson correlation coefficient of −0.27, showing that there is little to no correlation between the differential proteins identified by Mayer et al. and those identified by Johnston et al.

The lack of overlap in differential proteins between these two studies was worrisome, and we attempted to identify similarities in differential expression at the pathway level. We ran a KEGG pathway enrichment, separating the significantly upregulated and downregulated proteins for each study. We found that none of the pathways in the upregulated proteins were the same, and only Leukocyte transendothelial migration, Platelet activation, and Leishmaniasis were shared in the downregulated KEGG analysis (Supplementary Table 4). This further demonstrates that the differential proteins identified by Mayer et al.^14^ are not similar to those identified by Johnston et al.^13^ despite identifying many of the same overall proteins.

### Our study vs. meta-analysis

The goal of our study was to determine whether ultra-small sample proteomics could be used in longitudinal surveys of CLL patients. To test the efficacy of our ultra-small samples, we compared our total proteomic data to that of Mayer et al.^14^ and Johnston et al.^13^ We found that the majority of our proteins were identified by both papers (Figure 3), suggesting our data provides sufficient coverage for longitudinal CLL studies. Using the abundance values provided by Mayer et al.^14^, we discovered that our data identifies the many of the most abundant proteins in healthy individuals. We identified roughly 79% of the top quartile of proteins Mayer et al. identified, 30% of the second quartile, 10% of the third quartile, and 6% of the fourth quartile. Our proteomic data also identified roughly one third of the differential proteins identified by Mayer et al.^14^ (30.9%) and Johnston et al.^13^ (34.6%).

**Figure 3.**
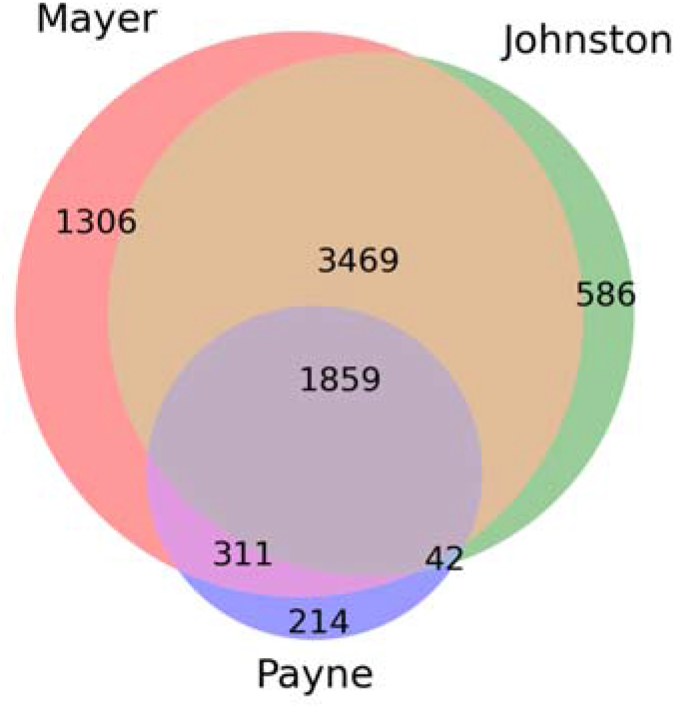
Venn diagram of total proteomic data from Mayer et al.,^14^ Johnston et al.,^13^ and our data. Within our GitHub repository (https://github.com/PayneLab/TwoMonthLymphocyte), the exact code for Figure 3 can be found in file “~/Make_Figure_3.ipynb”

## Discussion

Proteomic studies have already identified several protein markers for CLL as well as metabolic pathways that can be targeted with treatments^13,26^. However, such studies are often limited to single time point characterization. Our pilot study explored the temporal and individual variability of B and T cells from healthy subjects. We demonstrate that the proteomes of healthy individuals vary significantly between time points and between people. Since our dataset had five replicates per condition and high technical reproducibility (average correlation coefficient of 0.93), we are confident that the temporal variability we saw was true biological variability.

We performed a meta-analysis of two proteomics studies of CLL which showed substantial overlap in the proteins identified and little overlap between differentially expressed proteins. As seen in our results, the proteome of healthy individuals changes significantly with time. The lack of overlap in differentially expressed proteins identified by Mayer et al.^14^ and Johnston et al.^13^ could be caused by having only sampled at one time point and therefore not truly capturing a subject’s B-cell profile. Pathogen exposure, lifestyle changes or seasonal patterns could contribute to temporal proteomic changes^27^, suggesting that control subjects should be sampled at multiple time points.

An additional aspect of our pilot study is the proteomic identification of ultra-small samples (145 cells). Our analyses quantified 2426 proteins, and confidently identified canonical features of B and T cell function. The ability to analyze ultra-small samples allows for characterization of rare cell types and intra-tumor heterogeneity. Additionally, high quality proteome coverage of ultra-small samples enables more frequent sampling with lower blood volumes, which can be used to better understand leukemia evolution.

## Acknowledgements

This work was supported by the BYU Simmons Center for Cancer Research and NIH under award number R01GM138931.

